# A Multimodal Adaptive Super-Resolution and Confocal Microscope

**DOI:** 10.1101/397273

**Authors:** Liyana Valiya Peedikakkal, Andrew Furley, Ashley J. Cadby

## Abstract

Existing optical microscopy techniques compromise between resolution, photodamage, speed of acquisition and imaging in to deep samples. This often confines a technique to a certain biological system or process. We present a versatile imaging system which can switch between imaging modalities with sub millisecond transition times to adapt to the needs of a wide range of sample types. The imaging modalities provide the minimally invasive but low-resolution epi-fluorescence though increasing invasive but higher resolution confocal and structured illumination until the highest resolution is achieved through the most intrusive, localisation microscopy. The ability of the system to overcome the limitations of conventional single mode microscopy is demonstrated by several biological investigations. The ideas presented in this work allow researchers to move away from the model of a single imaging modality to study a specific process and instead follow those processes using the most suitable method available during the lifetime of the investigation.

---

Optical microscopy has played a significant role in biological research due to its non-contact and minimally invasive nature enabling *in vivo* investigation. Most imaging techniques are optimized for specific sample characteristics, for example, the ability to image deep into thick samples [1], to reduce the level of photo bleaching in samples [2], to image rapidly for live cell imaging [3] and most importantly the extent to which the technique can push the resolution limit [4,5,6]. Many super-resolution techniques such as Stimulated Emission Depletion (STED) microscopy [4] and Localization Microscopy (LM) [5,6] techniques apply high illumination powers and as a result can be damaging particularly to live samples. Low power and faster imaging give Structured Illumination Microscopy (SIM) [7] an advantage over other super resolution techniques. However, SIM is generally limited to resolution doubling.

The complexity and fragility of biological systems hinders the current state of the art imaging techniques from following live cellular processes, as a fixed modality is not suited to every time point of a sub-cellular phenomenon. The versatile imaging platform (VIP) presented here can adapt to meet the imaging requirements at specific time points within a sub-cellular phenomenon.

The mode of illumination and the mechanism by which the emitted light is detected are critical in determining image resolution and the extent of photodamage to the sample. In this work, we utilize a Digital Micro-mirror Device (DMD) based illumination and detection system to deliver the VIP which can switch between imaging modalities at sub-millisecond transition rates allowing us to combine low power widefield, structured illumination-based imaging and high-power single molecule localization microscopy into a single microscope.

By changing the direction of the DMD’s mirrors between two stable positions +12° and −12° which can be termed ‘ON’ and ‘OFF’ an illumination pattern can be projected on to a sample. When all the DMD pixels are ‘ON’, a uniform illumination is projected which can be used for widefield imaging and localisation microscopy. The average power of illumination can be controlled by rapidly switching between the two stable positions of the micromirrors at 22 kHz. Applying a pattern to the DMD allows us to perform a wide range of more complex imaging techniques such as confocal, SIM and LM (**Fig.1**).

To demonstrate the versatility of the system we present a number of biological investigations which could only be studied or are greatly enhanced by the adaptive nature of the imaging system. Firstly, we demonstrate the ability to switch between a low invasive and a higher resolution modality. We followed embryogenesis in drosophila through the initial nuclear divisions in the syncytial blastoderm stage of their development.

The adaptive nature of the system is used to address the fixed pinhole limitation of a conventional spinning disk microscope. The ability to change various confocal imaging parameters such as the pinhole size and the inter pinhole distance to suit a sample’s brightness and thickness is used to study thick auto-fluorescent samples. In the third example we show that using the efficient capture and control of light on the emission pathway allows us to apply a number of enhanced resolution techniques based on confocal microscopy. In the final section we demonstrate the application of epifluorescence to locate areas of interest for imaging followed by LM to image those areas with the highest resolution possible.

Drosophila melanogaster is a model organism widely employed to understand the development of single cells into complex multicellular organisms as well as the molecular basis of physiological mechanisms. Embryonic development in Drosophila is a rapid process where, after fertilization, a single cellular embryo develops into the cellular blastoderm consisting of 6,000-8,000 cells with a specific morphology and pattern formation which determines the future spatial organization of the fly’s body. Imaging techniques of various kinds have been employed in the study of this process. For example, electron microscopy has provided high resolution information at specific time points in fixed tissues [8], while spinning disk and light sheet microscopy have been used to image changes in living cells as development proceeds [9,10].

Among the critically important cellular processes occurring between zygote formation and the cellular blastoderm are the multiple mitotic nuclear divisions of the syncytial blastoderm, but it is known that light exposure or photodamage can slow down mitotic division limiting our ability to study this process in real time [11]. To study the dynamic cellular processes in live cell imaging with minimal photodamage, the VIP was used to change between the low invasive widefield modality to the more invasive, but higher resolution, confocal imaging at the start of a nuclear cycle. Here the dynamics of mitotic division are studied by imaging centrosomes which are the major microtubule organising centres of the cell, regulating cell-cycle progression.

Live Drosophila embryos were imaged using low light exposure minimal invasive widefield microscopy at the early mitotic division nuclear cycles. Figures 2(a), 2(b) and 2(c) are temporal snapshots of centrosomes at initial nuclear divisions during the development of the embryo. At the onset of the desired nuclear cycle of interest, just before the centrosomes split, the pattern on the DMD is switched and rapid confocal images are acquired until one cycle of division is completed **(Fig. 2(d)** and **Supplementary Video 1** and **3)**. Images are acquired using a 60x water objective for one cycle of centrosome division. The rapid confocal data can then be used with the Super Resolution Radial Fluctuation (SRRF) algorithm to provide high resolution information [12]. **Figure 2(e)** and **Supplementary Videos 2 and 4** show the high-resolution dynamics of centrosome splitting at specific time points in the syncytial blastoderm stage of the embryo development. Figure 2(d1-d5) and figure 2(e1-e5) are 5 time points of the region highlighted with white square from the of 100 time points confocal data (Fig. 2(d)) and reconstructed SRRF images (Fig 2(e)) respectively.

**Fig. 1.**
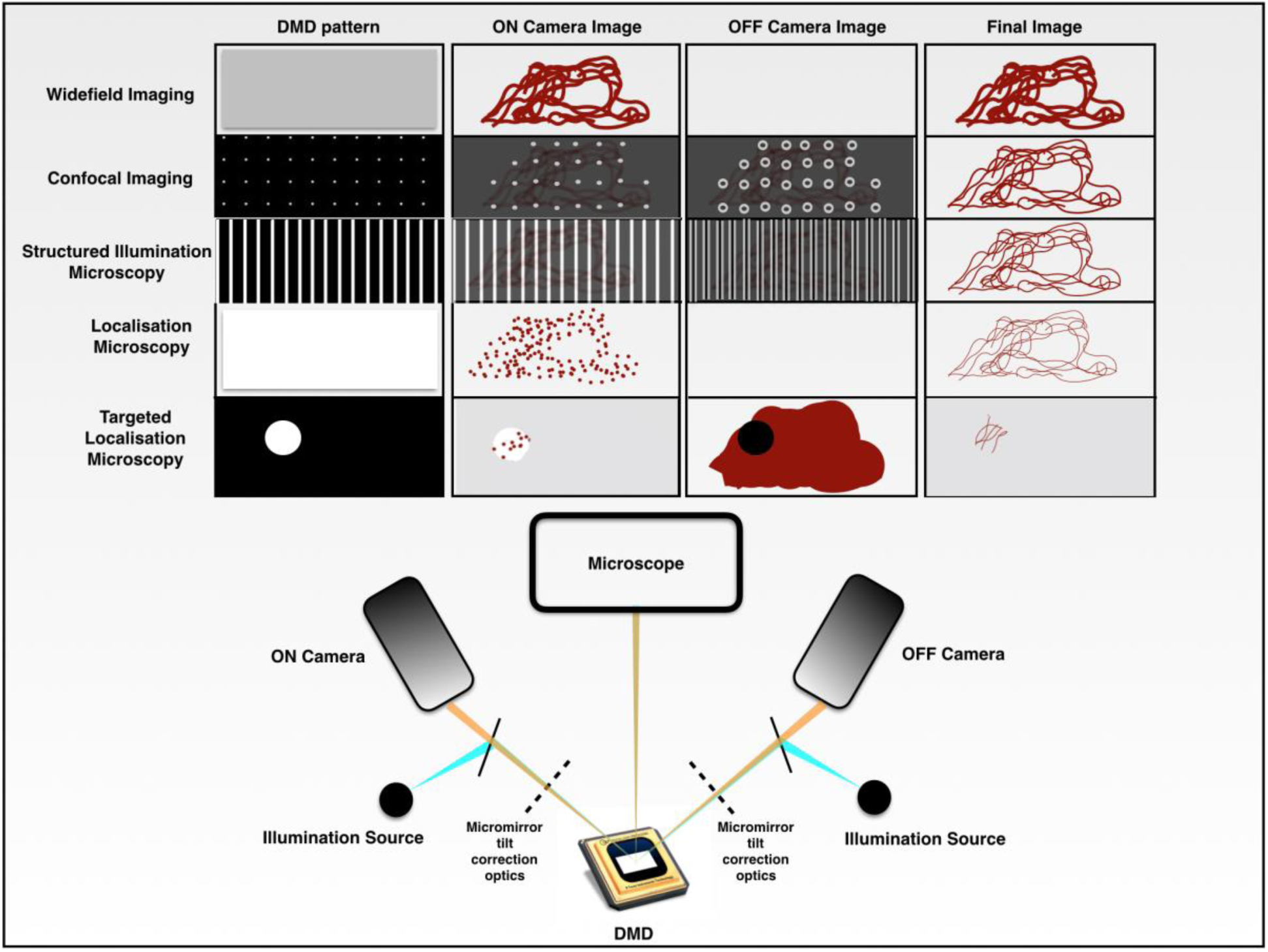
Key aspects for implimenting a multimodal adaptive super resolution confocal microscope. A DMD is used to project various patterns onto the sample, this allows for the development of the multimodal adaptive super resolution confocal microscope. Widefield and LM microscopy project a uniform pattern with low power and high power illumination intensities respectively. Conventional confocal microscopy projects a multifocal pattern and conventional structured illumination microscopy projects a lamellar pattern. The ON camera images illustrate a single frame acquired in the imaging modality. The OFF camera images illustrate a single frame collected in the imaging modality which is traditionally rejected by conventional optical setups. The setup can target specific regions of interest in the sample either for photomanipulations or as illustrated here for targted localisation microscopy. Final images (right) indicate that the control of power and pattern of illumination can tune resolution of the image and photodamge in the sample. Only the major components of the optical setup are given for clarity.

**Fig. 2.**
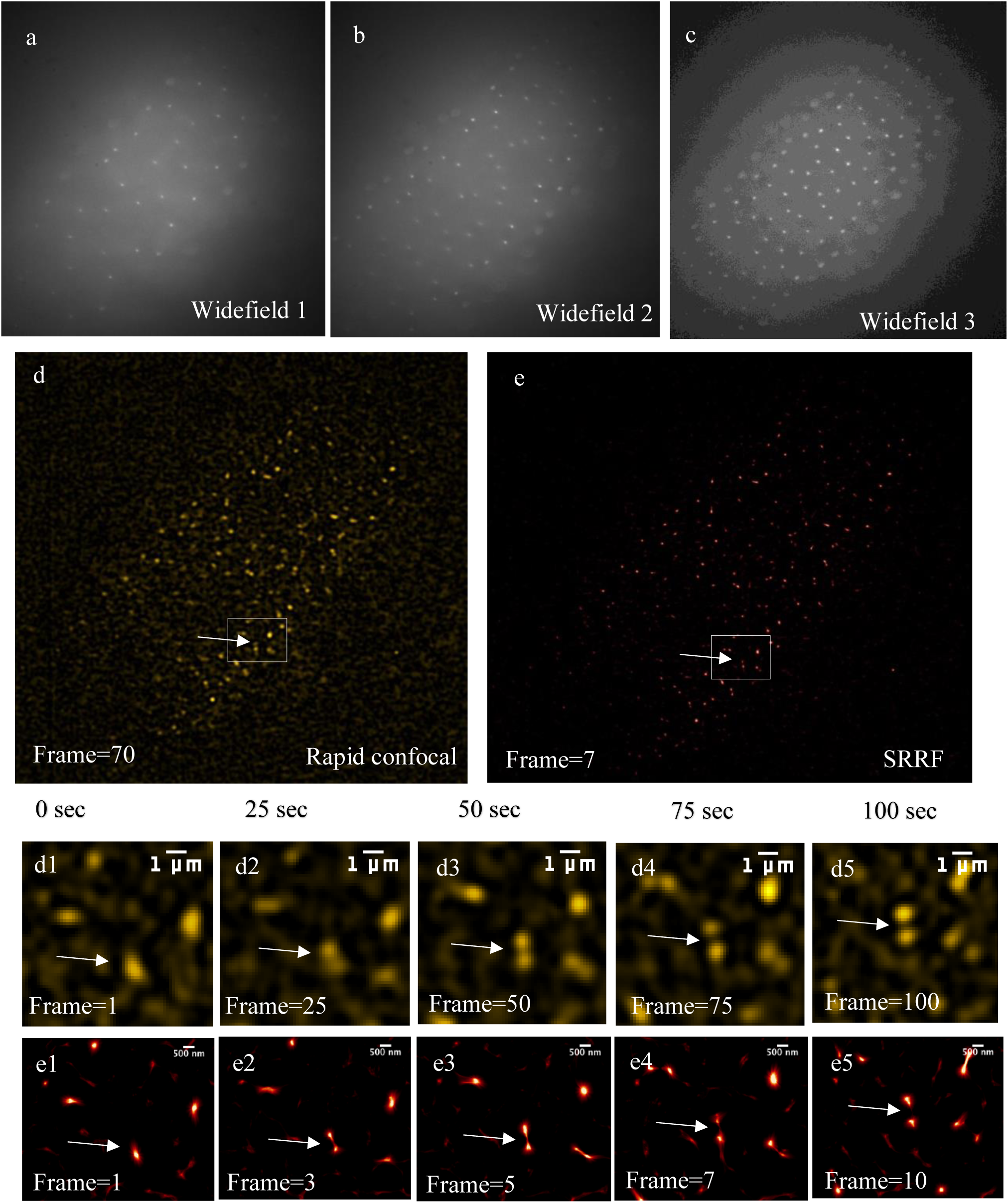
Early mitosis stage in drosophila embryogenesis in super resolution. (**a-c**) Widefield images of early mitotic divisions. Mitotic division is visualized with the division of centrosomes. (**d**) Frame 70 of Rapid Confocal. (**e**) Frame 7 of SRRF reconstruction of Rapid Confocal data. (**d1-d5**) Zoomed in Rapid confocal for a single centrosome splitting. (**e1-e5**) Zoomed in SRRF for a single centrosome splitting. **(d1-d5)** and **(e1-e5)** are frames at 0^th^, 25^th^, 50^th^, 75^th^, 100^th^ time points.

Changing between imaging modalities reduces the overall photodamage to the embryos. The switching of imaging modalities to visualize the dynamics of centrosome division provides higher resolution than previously visualized [13] and the versatility to rapidly switch between imaging modalities reduces photodamage during embryonic development. **Supplementary Figure 3** demonstrates the sub-millisecond transition times between imaging modalities.

**Fig. 3.**
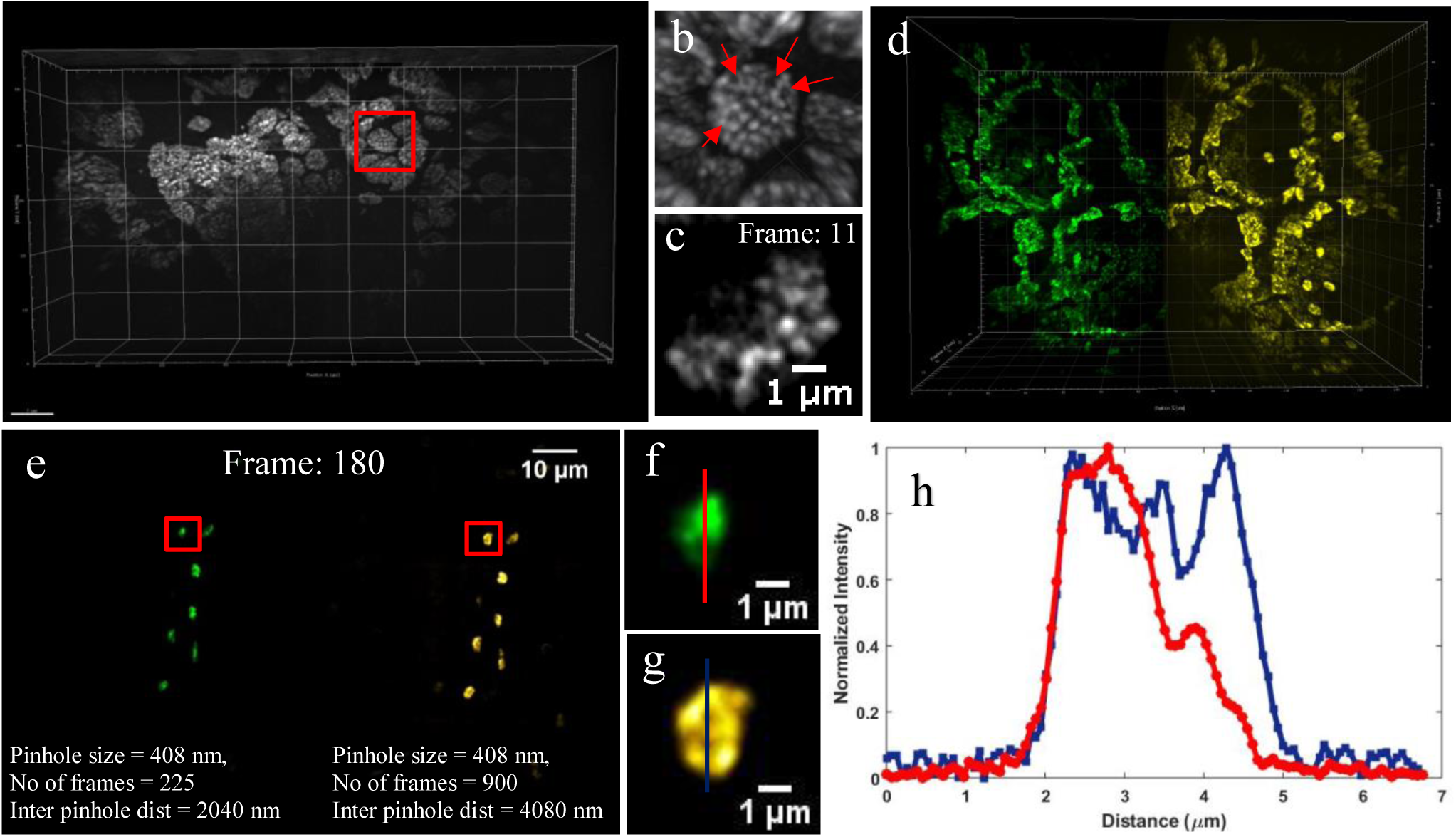
Adaptive confocal modality for deep imaging and revealing the grana and stroma lamellae of chloroplasts in situ in a leaf disk. (**a**) 30-degree perspective view of 3D Z-projection of rapid confocal for 50 slices with 15 µm depth. (**b**) Red square zoomed in, red arrows indicate the stroma lamellae of the chloroplast. (**c**) Frame 11 of the selected chloroplast of figure (**a**). This figure shows the position of grana in one slice of the chloroplast Z-stack image. Supplementary figure 7 shows the 23 slices Z-stack images through one selected chloroplast marked by red square in (a). 3D projection of 23 slice Z-stack of one chloroplast shows the location of stromal lamellae marked in red arrows in (b). (**d**) 30-degree perspective view 3D Z-projection of rapid confocal for 200 slices with 60 µm depth, green LUT is given for inter pinhole distance 2,040 nm and yellow LUT is given for inter pin hole distance 4,080 nm. (**e**) Frame 180 of figure (**d**). (**f-g**) Selected area zoomed in. (**h**) Plot of Normalized intensity of line profile in (**f**) and (**g**).) The rapid confocal raw data show above were deconvoluted using AutoQuant software with 10 iterations and minimum noise setting in adaptive PSF (blind) deconvolution.

The ability to rapidly switch between different confocal patterns is used to improve imaging quality in thick samples. In a conventional confocal system with constant inter pin hole distance, the intensity of out-of-focus light collected at the detector increases with imaging depth as the emission planes deeper in to the sample undergo the highest scattering when passing through the sample [14]. This increases the cross talk between the nearest pinholes and reduces image quality. Switching to a larger inter pinhole distance as the system images further in to the sample during acquisition is a unique advantage of the VIP.

**Figure 3** shows chloroplasts in *Spinacia oleracea* leaf disks imaged using the adaptive confocal modality of the VIP. Stromal lamellae are structures smaller than the diffraction limit, only a few nanometres thick as previously visualized using electron microscopy [15]. Due to their size and location it has been difficult to visualize these with SIM which has a limited axial depth or conventional confocal microscopy [16,17,18]. **Figure 3(a-c)** (**Supplementary video 5,6,7**) shows a 15 µm stack revealing the locations of grana in the chloroplasts. The Z-projection given in **figure 3(b)** of a single chloroplast, allows us to locate the stroma lamellae which are marked with red arrows. **Figure 3(b)** is the projection of 23 z-slices though a single chloroplast (full stack given in **supplementary figure 7)**, a single z slice (slice 11) is shown in **figure 3(c)**. With the single slice of the chloroplast the location of grana can be visualized (**Fig. 3(c))** and the 3D projection with 23 slices through one chloroplast (**Fig. 3(b))** shows the precise localisation of stromal lamella which are links between the grana in the chloroplast.

To demonstrate the improvement that can be achieved by altering the inter pin hole distances two datasets were collected deep into the sample (60 μm) using confocal patterns with different inter pinhole distances, 2,040 nm and 4,080 nm in the image plane. The data sets are shown as green and yellow respectively from now on and are presented in **figure 3(d)** as a projection through the sample. **Figure 3(e)** a single Z slice taken at 60 µm depth, shows several isolated chloroplasts. The areas highlighted by red boxes are given in **figure 3(f)** and **3(g)** clearly show the improved clarity obtained with the wider confocal pattern, with **figure 3(h)** showing the normalized intensity profile through the same chloroplast with two different inter pin hole distances. This allows us to resolve structures deep in to the sample when using a confocal pattern with a larger inter pinhole distance.

Although these structures have been seen with electron microscopes previously, we report the precise localization of stromal lamella in plant chloroplasts achieved with optical microscopy. The adaptive deep imaging modality presented in this work demonstrates the imaging of chloroplasts and stroma lamellae at 60 µm. This has, to the best of our knowledge, not been visualized previously at this clarity with an optical microscope.

Confocal microscopy is the most widely used microscopy technique for biological studies and has become the workhorse of biological imaging, but in its conventional form it remains diffraction limited. Obtaining super resolution images using confocal microscopy was first theoretically proposed in 1988 by Shephard [19] and experimentally achieved as Imaging Scanning Microscope (ISM) by Muller *et*. *al*. [20] in 2010. ISM has been implemented using various optical setups such as in a laser scanning confocal microscope [20], Digital Micromirror Device (DMD) in Multifocal Structured Illumination Microscopy (MSIM) [21], lenslet array in Instant Structured Illumination Microscopy (iSIM) [22], spinning disk in Confocal Spinning Disk-Image Scanning Microscopy (CSD-ISM) [23] and the AiryScan microscope [24].

In a conventional fluorescence microscope, almost 90% of captured fluorescence from a sample is out-of-focus [25]. One of the crucial elements when building a confocal based structured illumination microscope is the rejection of out-of-focus light. Despite these recent advances, implementations of ISM [20,21,22,23,24] do not efficiently utilize out-of-focus light because they are based either on physical pinholes which prevents the out-of-focus light from being detected at all (and thus this information is lost) or on digital pinholes which mix both in and out-of-focus light at ratios which vary throughout the sample, making utilization of information difficult. Using a DMD to provide both illumination and detection allows the collection and separation of light passing through the confocal pinhole and light that would normally be rejected by the pinhole [26–28].

Here we present super resolution adaptable microscopy (SR-AM), which improves the signal-to-noise ratio and resolution of the ISM implementation by efficiently using both in-focus and out-of-focus light from the sample. SR-AM is compared with conventional widefield and confocal microscopy (Fig. 4), and a recent software based ISM implementation **(Fig. 5).**

**Fig. 4.**
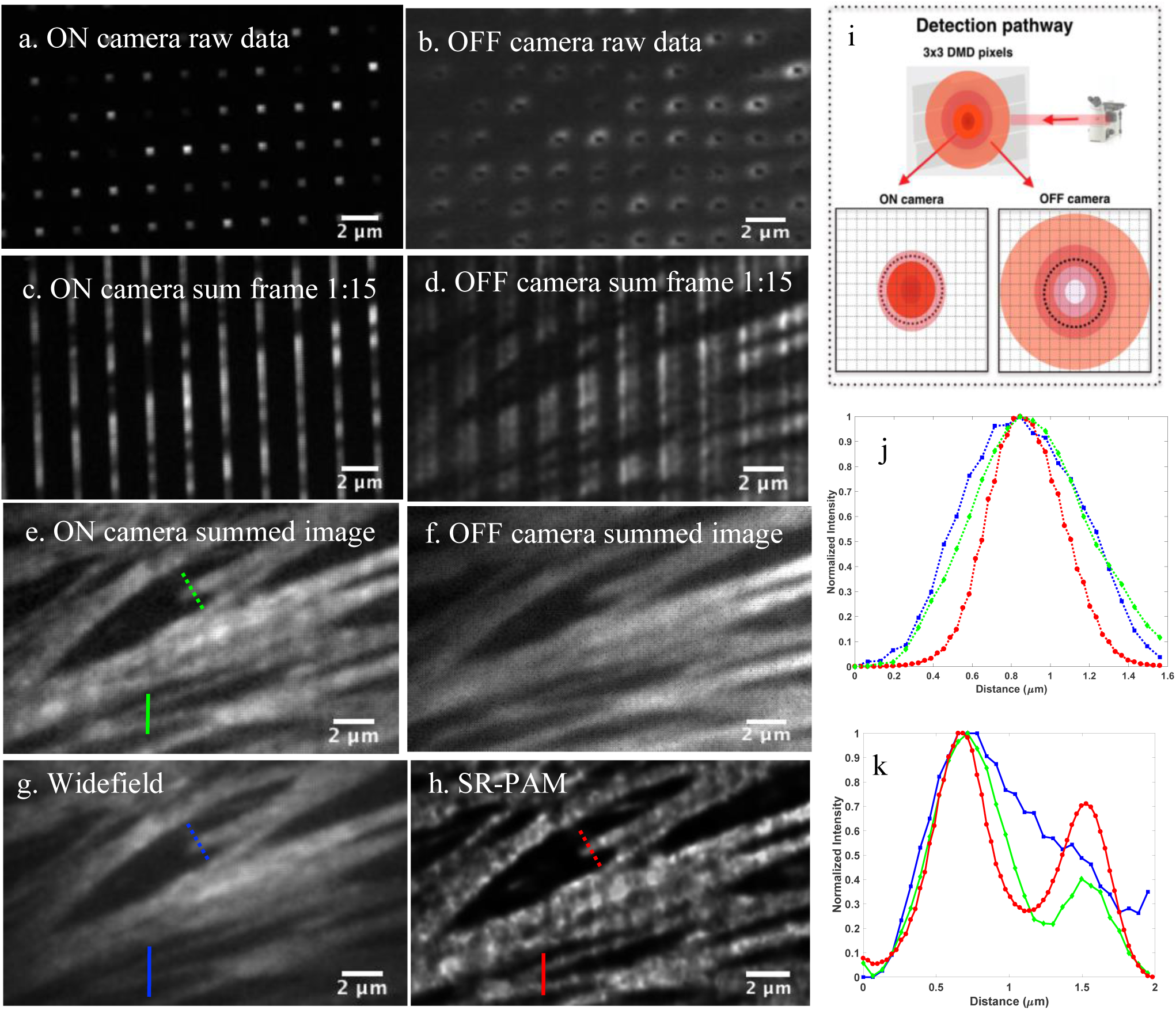
Bovine Pulmonary Artery Endothelial Cells (BPAEC) labelled for F-actin filaments with phalloidin are visualized to demonstrate the enhancement in contrast and resolution in SR-AM. (**a**) In-focus data collected in ON camera with 3×3 DMD pixels as pinhole size and 3×5 DMD pixels as distance between two pinholes. (**b**) Out-of-focus data collected in OFF camera. (**c**) Sum of frames 1 to 15 in ON camera. (**d**) Sum of frames 1 to 15 in OFF camera. (**e**) Sum of all the 225 frames in ON camera. (**f**) Sum of all the 225 frames in OFF camera. (**g**) Widefield image. (**h**) SR-AM image. (**i**) Optical pathway of detection of emission spots. (**j**) Plots of intensity along the respective dotted coloured lines in **(e), (g), (h)**, FWHM values are, widefield 780 nm, confocal 720 nm, SR-AM 460 nm. Plots of intensity along respective solid lines in **e, g, h.**

**Fig. 5.**
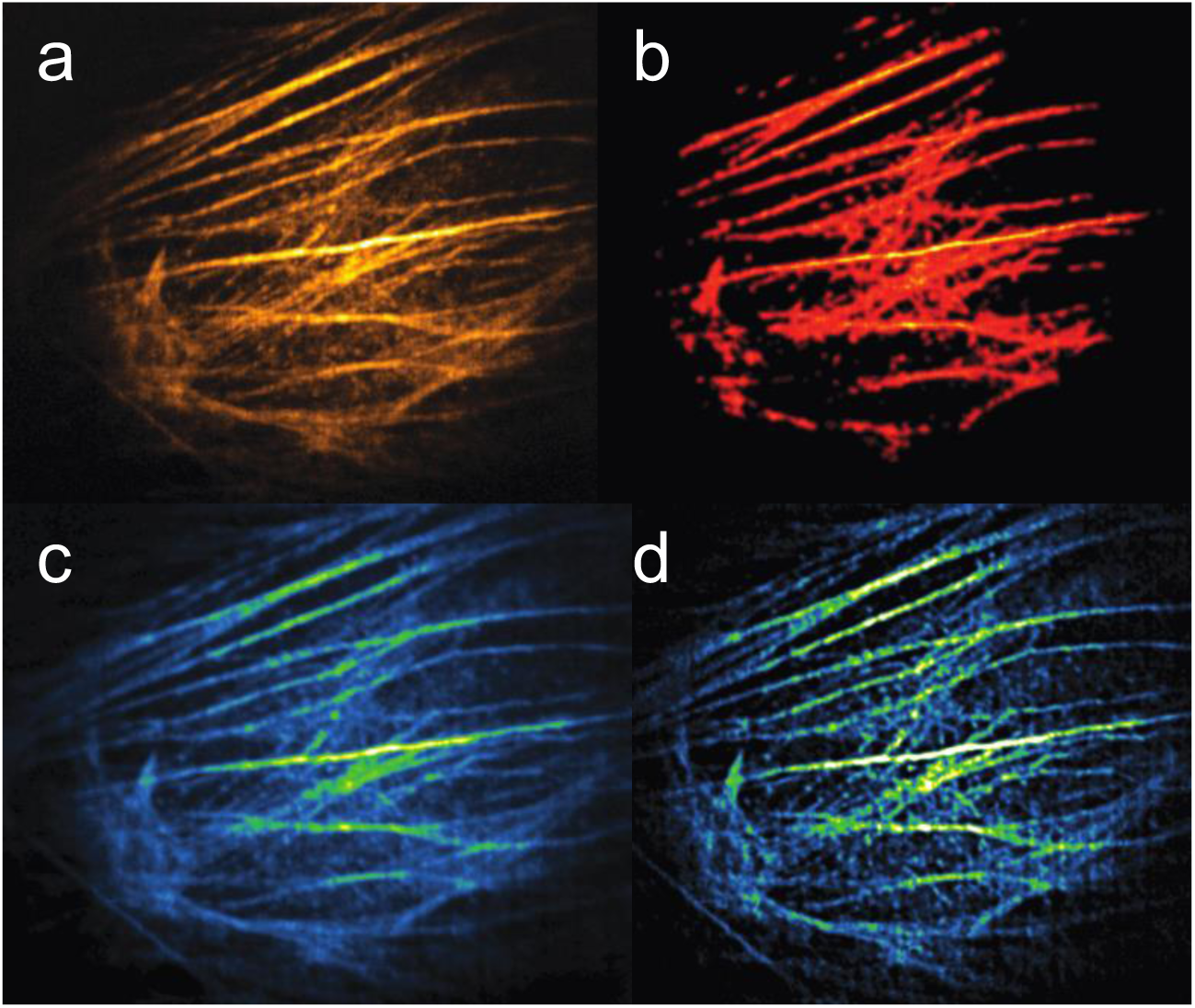
A comparison of confocal, CSD-ISM and SR-AM on BPAEC labelled for F-actin filaments with phalloidin. (a) The confocal image, data collected on the ON camera using the DMD as a physical pin-hole. This is a maximum value projection of the 225 frames collected, one frame for each DMD pattern. (b) The CSD-ISM image calculated on the single frames from part (a), the position of the confocal spots was calculated using Thunderstorm, this requires the selection of a threshold. (c) The SR-AM image is calculated using the methods outlined in this work. Mapping the DMD to the cameras allows for a background correction and the removal of threshold. (d) The SR-AM image after deconvolution with Huygens deconvolution with classic maximum likelihood deconvolution. Resolution enhancement after SR-AM post-processing depends on the deconvolution software applied.

In the VIP, in-focus light is collected by the ON camera, this is the same information as collected by a conventional laser scanning confocal microscope (LSCM) or spinning disk microscope and is shown in **(Fig. 4(a), 4(c), 4(e))**. The out-of-focus light from the sample is collected by the OFF camera **(Fig. 4(b), 4(d), 4(f))** which contains the information which is usually rejected by a conventional confocal microscope **(Supplementary video 8).** In the schematic for the optical pathway **figure 4(i)** one DMD pixel is turned ON (+12 degrees) and is surrounded by OFF (−12 degrees) pixels, the ON pixel directs the light towards the microscope. A single DMD pixel translates to 136 nm at the sample plane of the microscope. Light returning from the sample, after diffraction spreading, hits the ON pixel and the surrounding OFF pixels. The ON pixel acts as an analogue of a pinhole in a conventional confocal system and the in focus light from the sample is collected on the ON camera. The out of focus light collected in the surrounding OFF pixels, which would be rejected by a conventional confocal microscope, is directed towards the OFF camera, the black dotted circle on both cameras represents the size of a virtual pin hole in a conventional confocal system.

Using the data collected by the OFF camera a background correction of the image can be performed to improve the signal to noise ratio. For SR-AM, we subtract a portion the out-of-focus Point Spread Function (PSF) from the in-focus PSF to remove background light and to achieve a higher signal-to-noise ratio. This makes efficient use of all of the collected light from the sample (**supplementary note 1**). The out-of-focus PSF subtraction method **(Online Methods** and **Supplementary note 6)** is given in the supplementary materials **(Supplementary note 2)** as well as simulations **(Supplementary note 3)**.

The resulting corrected PSF is then used along with a Sheppard summing [19] algorithm to increase the resolution. When a low frequency multifocal illumination pattern mixes with the sample frequency, higher frequency sample information is collected by frequency mixing at the objective. Reducing the size of the collection pinhole in the image plane increases the collection of this high frequency information from the objective’s Fourier plane. To extract the high frequency information a software based displaced point detector technique is employed to collect all the emission intensity without the loss of signal due to reduced size of pinhole. This is followed by pixel reassignment to bring the intensity of the displaced pinhole back to the optical axis to enhance the resolution of the image. The SR-AM theory is an extension of the conventional programable array microscopy [29] and ISM [19] and is provided in **supplementary note 2.** The resolution enhancement achieved using Shepard summing is shown for an F-actin labelled BPAEC cell in **fig. 4(h)** and compared with the widefield **(Fig. 4(e))** and the confocal image **(Fig. 4(g))**. Line profiles through the confocal image, the wide field image and the SR-AM image are given in **figures 4(j)** and **4(k)**. These show an increase in resolution and much higher signal to noise.

We now compare our implementation of ISM, SR-AM, with conventional confocal microscopy and a software-based ISM to demonstrate the superior background rejection afforded by using the DMD in both the collection and excitation pathways. The ISM compared in this work is the CSD-ISM implementation which has applied resolution enhancement with a spinning disk optical setup using a stroboscopic method to capture individual frames [23].

In the VIP’s optical setup, data collected on the ON camera is equivalent to a conventional confocal system with the DMD micromirrors acting as an analogue of a conventional confocal aperture. Imaging an F-actin labelled BPAEC the conventional confocal image is collected using the ON camera and given in **(Fig. 5(a))**. This confocal data can then be used to obtain the software-based ISM image. The data is processed using the CSD-ISM software to obtain and ISM image **(Fig. 5(b))**. The ISM processing used in **Fig. 5(b)** requires an initial step to identify the confocal spots using localization software such as ThunderSTORM [30] or RapidSTORM [31]. This localization requires a threshold, which can be used to limit the out-of-focus light and is discussed in **supplementary figure 9**. The thresholding method can be inefficient as confocal spots with low intensity can be missed. SR-AM does not require any thresholding as emission spot finding is based on calibrating the exact expected emission spot pixels in both ON and OFF cameras. This is achieved by precisely mapping the DMD pixels with both the ON and OFF camera pixels. Our DMD-camera calibration technique which achieves a precision with an error of less than 130 nm (1 DMD pixel) is given in **supplementary note 4**. The resultant background corrected SR-AM image can be seen in **Fig. 5(c)** along with a deconvoluted image in **Fig. 5(c).**

The VIP’s optical setup also allows us to obtain MSIM and AiryScan equivalent images which are based on collecting the full emission spot without physical apertures to reject out-of-focus light, this is achieved by adding the ON and OFF emission spots collected on the two cameras to restore the full PSF followed by employing MSIM and AiryScan (**Supplementary note 7)** equivalent algorithms (provided with this paper). A resolution comparison of SR-AM is performed on 100 nm bead samples and compared with resolutions achieved using widefield imaging, confocal and MSIM **(Supplementary note 1)**.

To establish the ability of the VIP to adapt to biological investigations requiring higher resolution, localisation microscopy (LM) is implemented. NIH 3T3 cells, shown in **figure 6**, were imaged using a combination of widefield and Stochastic Optical Reconstruction Microscopy (STORM) imaging to investigate the subcellular localization of a neural cell adhesion protein, NrCAM [32]. Specifically to test whether it can be found in primary cilia, a microtubule-based cell surface projection. NrCAM, transiently expressed by NIH3T3 cells after transfection, is labelled with Alexa Fluor 647-labelled secondary antibodies. The cilia are visualized with rhodamine red-labelled antibodies to the cilium-specific Arl13b protein [33].

**Fig. 6.**
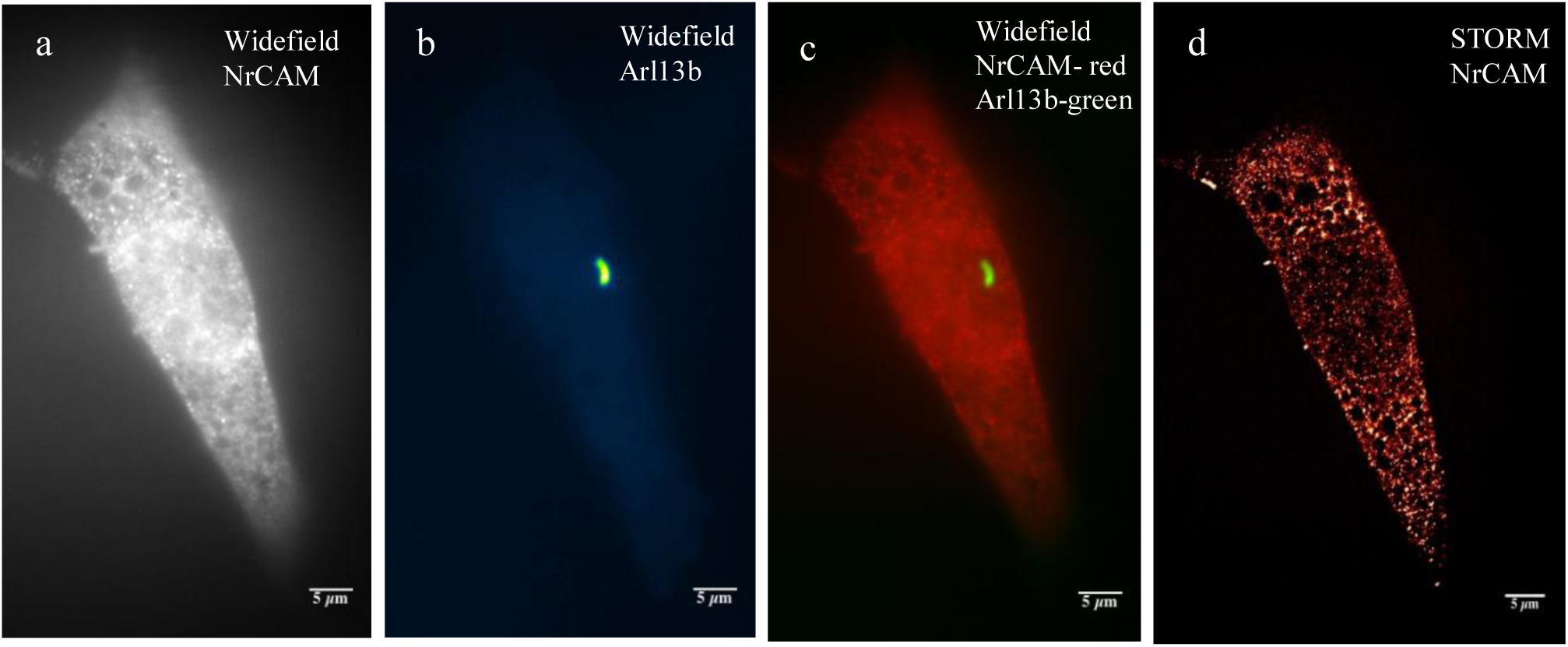
NIH 3t3 cells expressing NrCAM and Arl13b in widefield and STORM for super resolution localization of proteins at specific locations in the cell. (**a**) Widefield image of NrCAM with low power 647 laser in optical pathway 1 of the SR-AM setup. (**b**) Widefield image of Arl13b in cilia with 532 LED illuminated from the optical pathway 2 of SR-AM setup. (**c**) Merged image of (**a**) and (**b**) to locate the position of cilia with respect to the localization of NrCAM in the cell. (**d**) Super resolution localization of NrCAM protein in the cell implementing STORM microscopy with high power 647 laser illumination from the optical pathway 1.

The two independent optical pathways in the VIP are used, one to deliver low intensity light and one for the high-power laser illumination required for LM. This system can switch between the two pathways, light sources, imaging techniques, illumination wavelength and intensity, at sub millisecond timescales. Here the Arl13b is used to locate the position of cilia within the cell using low phototoxicity widefield illumination. Under low light widefield imaging the cell is scanned in the z direction to locate the cilia and then excitation pathway is switched, by changing the DMD pattern, to the path containing the high power 647 nm STORM illumination.

The VIP with super resolution localization reveals the presence of NrCAM on what appears to be the vesicular transport system of the cellular cytoplasm but indicates that there is no substantial NrCAM localization in the cilium. Unlike conventional LM in which the entire sample is illuminated with high laser power, we are not constrained to exciting the whole field of view. The versatility of this system to allows selective illumination of sections of interest in order to achieve targeted super resolution images. **Supplementary note 9** shows further examples of targeted illumination, where single bacteria of interest in a sample are selectively illuminated.

With the calibration code for the high precision for mapping DMD to camera pixels **(Supplementary note 4)** and the software for targeted illumination **(Supplementary note 9),** we provide a tool which can be easily integrated and widely adapted for optogenetics [34], Florescence Recovery After Photobleaching (FRAP) [35], photoactivation [36], ablation [37], photo-conversion [38] and targeted LM.

A pathway to a versatile multimodal imaging platform is presented here which allows biological processes to be followed through a range of imaging techniques. The system can rapidly switch between the minimally invasive but low-resolution epi-fluorescence though increasing invasive but higher resolution confocal and SR-AM until the highest resolution is achieved through the most intrusive, localisation microscopy. The novel SR-AM modality outlined here achieves a two-fold resolution improvement in a manner similar to previous implementations of SIM such as MSIM, instant SIM, and CSD-ISM. SR-AM improves on these previous techniques by utilising the emission intensity of both the in and out of focus light to improve the resolution and contrast. For the highest resolution achievable localisation microscopy in the form of STORM can be applied to the sample, this effectively allows the user to tune resolution as a function of photo-toxicity.

This has allowed us to follow a live process in drosophila under a minimally invasive wide imaging modality and seamlessly switch to confocal to achieve higher resolution images with improved background rejection when required. The adaptive nature of the system allows us not only to change the imaging modality but also to adapt a modality in real time to suit the sample. The versatile imaging modality presented here with adaptive confocal has the ability to change the pinhole size and the inter pinhole distance in real time during image acquisition implementing adaptive confocal for deep imaging reducing cross-talk at larger sample depths. A background corrected confocal spot allows us to perform a number of super-resolution methods and algorithms such as the AiryScan method.

This allows researchers to move away from the model of a single imaging modality to study a specific process and instead follow those processes using the most suitable method available, adapting during the lifetime of the investigation.

## Acknowledgements

We thank Imene Bouhle (University of Cambridge) of the Conduit lab for drosophila sample preparation and Sam Barnnett (University of Sheffield) of the Hunter lab for the leaf disk sample preparation. We thank Jörg Enderlein (Georg-August University, Göttingen) for providing us with CSD-ISM code. E.coli targeted illumination system was developed and tested on a mother machine system from the Jun lab (University of California, San Diego). The VIP and SR-AM was implemented on a prototype designed by Martin Thomas (Cairn Research). L.V.P is grateful for the PhD studentship through the University of Sheffield funded 2022 Futures scheme Imagine: Imaging Life. The authors would also like Ricardo Henriques (University College London), Jamie Hobbes (University of Sheffield) and Cairn Research (Faversham, UK) for critical reading of the manuscript. This work is supported by the Imagine: Imaging Life initiative at the University of Sheffield.

## Author Contributions

L.V.P and A.J.C conceived the idea and designed the experiments. L.V.P conducted the experiments and collected the data. L.V.P and A.J.C built all the software, analysed the data and wrote the manuscript. A. F supervised NIH 3t3 experiments and helped with manuscript writing. A.J.C designed and supervised the research.

## ONLINE METHODS

### Instrumentation of SR-AM

All optics were fixed in a 195 cm x 120 cm x 80 cm optical table (Thor Labs) to minimize mechanical vibrations. Nikon Eclipse Ti was used as the microscope base which held objectives (100x and 60x). Z-stage piezo (Prior Nano Scan Z) was used to translate the sample in the Z direction. Optical setup was relayed to the side port of the microscope. A Photometrics 95B is attached to the backport of the microscope (this is optional) which could be used to collect data without physical pinholing or the emission optical pathway passing through the DMD and could be used for comparison. Two filter cube holders were kept empty when imaged using the cameras attached with the DMD optical setup or an appropriate dichroic mirror or beam splitter could be used to image with the camera at the back port of the microscope. An Obis coherent 488 laser, coherent sapphire 514 and coherent sapphire 561 lasers were relayed to one illumination path of SR-AM system and led light source (Cairn OptoLED) was attached to the second illumination path which allowed easy swapping between laser and LED illumination. Two illuminators could be used simultaneously in the two independent pathways of the optical setup. Illumination light after passing through excitation filter and dichroic filter, passes through a relay optics consisting of a convex mirror (focal length = −100 mm, AR coating between 425 nm to 875 nm), concave mirror (focal length = +250 mm, AR coating between 425 nm to 875 nm). These two mirrors pairs were collectively used in a Schwarzschild configuration, with an effective focal length of 165 mm. Two fold mirrors were incorporated between the concave mirror and the DMD in order to make the system more compact, and all mirror profiles were within L/4 per 25mm. For illumination, light was introduced into the infinity space prior to the convex mirror, offset from the Schwarzschild optical axis by a distance so as to arrive at the DMD at an angle of 24 degrees. Light falling on pixels tilted by 12 degrees in this same direction was thereby orthogonally reflected and relayed to the microscope objective. For confocal detection, light from the microscope followed the reverse pathway through the Schwarzschild mirrors and was then focussed to form the camera image by conventional lens optics. Imaging light falling on the OFF pixels of the DMD was similarly collected and sent to a second camera by the other Schwarzschild mirror pair. The refocussing lenses were similarly offset from their Schwarzschild optical axes, so as to form standard orthogonally focussed images at the camera, at or close to diffraction-limited quality [39], rather than the 24 degree tilted focus produced by previous DMD-based confocal systems, and which has discouraged their more general use.

Texas Instruments DLP7000 chipset with FPGA controller was purchased from Vialux. DMD is a bi-stable device with around 8 million micromirrors mounted on a CMOS memory cell arranged in 768 rows and 1,024 columns. The second plane fold mirror which reflects illumination light to the DMD and collects emission light from the DMD was aligned at +/-24 degrees for ON and OFF optical pathways respectively. DMD micromirrors have two stable positions +/-12 and the orientation of these micromirrors defines the DMD pattern. Custom made MATLAB GUI was used to generate and load the desired patterns on the DMD. For confocal imaging, multifocal patterns were generated to increase the speed of imaging and for widefield or STORM imaging, a uniform illumination pattern was used. Codes for projecting various patterns using DMD and all the patterns used in this paper in the format which are ready to be loaded on, to the DMD are available in supplementary code. Light from the illumination pathways forms a pattern at the DMD and is reflected along the optical axis perpendicular to the DMD face. Light from the DMD is collected and relayed to the side port of the microscope using two lenses (triplet lens with focal length 160 mm). There was a 1:1 magnification of beam between DMD to microscope side port. Light beam was de-magnified using a 100x oil TIRF objective before illuminating the sample.

Emission light coming back from the sample was collected by the same objective, passing through the two relay lenses and projected onto the DMD. In confocal imaging, when multifocal patterns were projected, each DMD micromirror acts as a physical pinhole to direct the in-focus and out-of-focus light from the sample into two pathways in the system. Emission light from the in-focus plane of the microscope follows the same pathway as the illumination light and was reflected to the ON pathway while out-of-focus light was reflected to OFF pathway and was collected in separate ON and OFF cameras respectively. Andor Zyla 4.2 sCMOS cameras which has a 6.5 µm pixel size were used for detection in the pathways. In the ON pathway, illumination beam and emission beam were separated at the dichroic. The emission beam passing through the dichroic is collected and focused to the camera sensor by lenses (triplet lens with focal length 160 mm) and a plane mirror. A detailed layout of optics and ray diagram of the optical setup is illustrated in **supplementary figure 1 and supplementary figure 2**.

### Data acquisition for confocal

For confocal imaging, this highly versatile system can operate in three modalities with rapid confocal which can run at 355.11 confocal scans per second, SR-AM which has a more revolutionary out-of-focus subtraction mechanism and adaptive confocal to change inter pinhole size and distance for deep imaging. For all imaging modalities except SR-AM, camera and DMD are running in its internally defined timings which allow us to change the camera exposure time depending on the sample brightness. For SR-AM, NI DAQ (USB-6341) was used to trigger the Z stage and ON camera. In this setting, ON camera is configured to trigger the DMD frame change and OFF camera exposure time as described in **supplementary figure 4**. It is worth noting here that these triggering settings are very crucial to get each camera frame to be in perfect sync with DMD frame changes achieving high level precision and control over each in-focus and out-of-focus emission spot. Z stage moves to the next axial level only after projecting all the confocal frames in one sequence.

### Speed of rapid confocal imaging

For rapid confocal imaging, the DMD was configured in the optical setup to act as a physical pinhole to reject the out-of-focus light rapidly to achieve high speed. In this mode, DMD is set to run at the fastest possible frame rate which is 44 µsec per frame. Modulation patterns employed here are multi-spot frames which are scanned to project point based illumination in sample plane **(Supplementary note 5).** Using a 100 frame confocal pattern, it acquires one confocal image in 4.4 msec. The speed of imaging is limited only by the speed of the camera and the brightness of the sample. When running the system in rapid confocal mode, the ON camera collects data which are equivalent to that collected in a spinning disk setup. The speed of the DMD allows us to image at frame rates of 355.11 confocal scans per second (when using 64 frames at 2×2 DMD pixels as pinhole size and 2×4 DMD pixels separation between pinholes). Confocal scans per second can be varied depending on the distance between the pinholes and the size of the pinholes. For 3×3 pinhole size, it can be varied from 101.01 (when using 225 frames at 3×3 DMD pixels as pinhole size and 3×5 DMD pixels separation between pinholes) to 280.58 (when using 81 frames at 3×3 DMD pixels as pinhole size and 3×3 DMD pixels separation between pinholes). 2×2 confocal pinhole which projects a 272 nm spot on the sample plane is the ideal pinhole size (**Supplementary figure 8**). We also use 3×3 pinhole size to deliver higher illumination intensities for a less bright sample.

### Data processing

In all optical setups with point based illumination and a physical pinhole based out-of-focus light rejection such as in LSCM, spinning disk, iSIM etc, there is an intensity blead through from the out-of-focus planes which is always collected in the detector. Other optical setups which are based on digital pinholing to subtract out-of-focus light by multiplying the full emission spot with a Gaussian mask, such as in MSIM, the out-of-focus blead through intensity is not subtracted. Moreover, out-of-focus intensity change throughout the sample due to the high inhomogeneous nature of biological structures which makes digital pinholing systems not fully efficient. **Supplementary video 8** shows raw confocal pattern scanning and reconstruction in ON and OFF camera. It can be seen from the video that out-of-focus light intensity varies for each illumination spot. SR-AM collects both in-focus and out-of-focus light in two separate cameras and calculates the intensity blead through from the collected OFF camera data during post processing and subtracts this from the ON camera data. Post processing is performed with custom software, written in the MATLAB programming language. Post processing includes: (i) two-tier DMD calibration with the ON and OFF cameras to achieve not more than two camera pixels error or 130 nm precision mapping between DMD and camera pixels, (ii) weighted subtraction of blead through intensity from each ON emission spot using the corresponding OFF emission spot to get the accurate PSF. (iii) pixel reassignment of the accurate PSF to account for ISM with v2 improvement in resolution and stitching all the spots to construct the final image and (iv) deconvolution of the final image to double the resolution. There are many DMD and camera-based techniques developed recently in optogenetics and particle tacking for example, and these need a high level of calibration between DMD pixels with the corresponding camera pixels to be effective. SR-AM calibration code can be adapted even for a non-confocal work which requires DMD and camera calibration and achieves less than 130 nm precision (which is the effective DMD pixel size at the image plane), we report in this paper. We provide the SR-AM reconstruction codes in supplementary code.

For versatile imaging, imaging modalities are switched instantaneously in software as demonstrated by switching between confocal and widefield in **supplementary figure 3**. STORM images acquired on the in-focus camera for **figure 6** is reconstructed using ThunderSTORM ImageJ plugin. Rapid confocal does not require any specific post processing as the image acquired in the camera is physically pinholed at the DMD to reject the out-of-focus light. Optionally, deconvolution and super resolution reconstructions can be employed as discussed in the next section.

### Deconvolution and super resolution reconstructions

Fast confocal data can be run through a deconvolution algorithm to restore the image and improve the resolution. We use AutoQuant (Media Cybernetics) with 10 iterations using adaptive PSF (Blind) method for data in **figure 3**. For rapid confocal imaging in **figure 2**, the final data were deconvoluted using Huygens deconvolution (SVI) with Classic Maximum Likelihood Estimation which improved the quality and restored the final image. Even though our detection of confocal emission light is based on DMD with micromirrors acting as physical pinholes, with the unavoidable loss of higher diffraction orders, we get enough signal-to-noise ratio to run the data through super resolution reconstruction algorithms such as 3B, SOFI etc, with the most recent Super Resolution Radiant Fluctuations (SRRF) algorithm to get super resolution images from the rapid confocal imaging mode.

### Adaptive confocal

A leaf disk is imaged in rapid confocal mode with a 100x oil 1.49 NA TIRF objective, 488 laser illumination and the emission light are collected using a 488/562 dichroic with a dual band FITC/Cy3 collection filter. Custom made software for pattern projection provided with this paper can change between confocal imaging with different pin hole sizes and inter pin hole distances. When imaging thick samples in an epi-fluorescence arrangement, imaging near the coverslip may not suffer from scattered light but imaging deep into the sample can increase the scattering and crosstalk between the pinholes degrading image quality. A smaller inter pinhole distance near the coverslip and larger inter pinhole distance deep inside the sample can reduce the scattering effect without any substantial decrease in image resolution. Adaptive confocal can either run in rapid confocal mode or with superior background correction and resolution doubling using SR-AM mode, depending on the sample characteristics and imaging requirements.

### Comparative MSIM and AiryScan processing

Multifocal Structured Illumination Microscopy (MSIM) is based on collecting the full PSF and digitally pinholing to reject the out-of-focus light, followed by Shephard summing and deconvolution to double the resolution. We can obtain an MSIM equivalent image using the data collected in our SR-AM optical setup by adding the ON and OFF emission spots to restore the full PSF and then following MSIM reconstruction steps. Resolution obtained with SR-AM is compared with MSIM in **supplementary note 1**. SR-AM enhances the signal to noise ratio of the image with its superior out-of-focus rejection mechanism and improvement in the resolution can be attributed to this. AiryScan microscopy which uses GaAsP detector obtains 32 raw images from 32 GaAsP detectors arranged to collect the full PSF. AiryScan reconstruction involves independent Wiener filter deconvolutions of the 32 raw images followed by Shephard summing. We can obtain an AiryScan equivalent image with the data taken in SR-AM optical setup by replacing the GaAsP detector with an sCMOS camera detector. The method detailed in **supplementary note 7** can be used with any confocal setup with a camera detector to obtain Airy images. We term this method as ‘AiryImaging’. MSIM and AiryScan equivalent codes for SR-AM optical setup are also provided with this paper in supplementary code.

### Plant sample preparation

Leaf tissue was prepared from market bought spinach leaves. A small leaf disk was placed on a microscope slide and mounted in vectashield (Vectalabs) under a.15 mm thick coverslip. **Samples in figure 4.** Bovine Pulmonary Artery Endothelial Cells (BPAEC) labelled with Texas Red-X phalloidin for F-actin and anti—bovine α-tubulin mouse monoclonal 236-10501 in conjunction with BODIPY FL goat anti—mouse IgG antibody labelling microtubules are used. Prepared slide was purchased from Thermofishers (product number-F14781). **Samples in figure 5**. F-actin was stained with Alexa Fluor 488 phalloidin. (Thermofishers, product number-F36924).

### NIH 3t3 sample preparation

NIH 3t3-L (3t3) mouse embryonic fibroblast cells were cultured with routine cell culture procedures on 22×22 mm size and 0.13 mm - 0.17 mm thick high precision coverslips. Coverslips were cleaned in 1M HCl for 30 minutes followed by cleaning in distilled water for three times and were stored in 100% ethanol for localization microscopy experiments.

Cells are grown in growth medium (10%FBS+Pen/Strep+DMEM) at 37 °C incubator). They need to be serum starved to allow for the growth of primary cilia. Primary cilia are present on dividing cells in only one phase of the cell cycle (G1), but this is very transient in rapidly dividing cells. Serum starvation makes the cells quiescent, increasing the number of cells in G1 and so increasing the proportion of cells bearing cilia [33]. Growth medium with 0.5% FBS in DMEM which contains 1% Glutamine and 1% Penicillin-Streptomycin was added to the cells and kept in incubatror overnight. Cells were prepared to express NrCAM protein through transfection. For transfection of NrCAM-HA plasmid was introduced using Lipofectamine-2000 (11668019, Invitrogen) according to supplier’s recommendations. Addition of plasmid ensures that protein we are intending to look at is expressed. 500 ng of NrCAM is added to each well. The cells were fixed the following day (after 18-24 h at 37°C) using 4% Paraformaldehyde (PFA) in 0.1M phosphate buffer at pH 7. The wells are placed on rocker for 20 minutes. PFA fix was removed and the wells were placed in rocker for 5 minutes in PBST (Phosphate Buffer Saline and 0.05% Triton). 3% normal goat serum was used as blocking serum. Blocking serum is added to cells for 1 h before immunolabelling [32].

Immunolabelling was performed as NrCAM tagged with HA was detected by anti-HA rat monoclonal (11867423001, Sigma Aldrich; 1:1000 in PBS, 0.05%TX100, 3% normal goat serum (PBST/NGS), incubated overnight at 4°C), which in turn was detected by Alexa Fluor 647 Goat Anti-Rat IgG (A21247 - Thermo Fisher Scientific) secondary antibody (1:200 in PBST/NGS, incubated 1-2 h in dark at room temp). Arl13b (11867423001, Sigma Aldrich; 1:1,000 in PBS, 0.05%TX100 and 3% normal goat serum (PBST/NGS) were used as primary antibody to label micro cilia. Rhodhamine red was used as secondary antibody. For STORM imaging, coverslips need to be mounted over 2 or 3 drops of STORM buffer. STORM buffer for this work was prepared by mixing 100 millimolar monoethanolamine (MEA), 0.5 mg/ml glucosidase, 40 μg/ml catalase and 10% w/v glucose in PBS solution [40].

